# An early-morning flowering trait in rice helps retain grain yield under heat stress field conditions at flowering stage

**DOI:** 10.1101/2021.01.16.426929

**Authors:** Tsutomu Ishimaru, Khin Thandar Hlaing, Ye Min Oo, Tin Mg Lwin, Kazuhiro Sasaki, Patrick D. Lumanglas, Eliza-Vie M. Simon, Tin Tin Myint, Aris Hairmansis, Untung Susanto, Bharathi Ayyenar, Raveendran Muthurajan, Hideyuki Hirabayashi, Yoshimichi Fukuta, Kazuhiro Kobayasi, Tsutomu Matsui, Mayumi Yoshimoto, Than Myint Htun

**Affiliations:** Japan International Research Center for Agricultural Sciences (JIRCAS), 1-1 Ohwashi, Tsukuba, Ibaraki 305-8686, Japan; International Rice Research Institute (IRRI), DAPO Box 7777, Metro Manila, the Philippines; Hokuriku Research Station, Central Region Agricultural Research Center, National Agriculture and Food Research Organization (CARC/NARO), Inada, Joetsu, Niigata 941-0193, Japan; Division of New Genetics, Yezin Agricultural University (YAU), Nay Pyi Taw 15013, Myanmar; Department of Agricultural Research (DAR), Yezin, Nay Pyi Taw 095-067, Myanmar; Indonesian Center for Rice Research (ICRR), Street 9th Sukamandi, Subang, West Java, Indonesia; Department of Plant Biotechnology, Tamil Nadu Agricultural University, Coimbatore 641003, India; Institute of Crop Science, NARO, 2-1-18 Kannondai, Tsukuba, Ibaraki 305-8518, Japan; Tropical Agriculture Research Front (TARF), Japan International Research Center for Agricultural Sciences (JIRCAS), 1091-1 Maezato-Kawabaru, Ishigaki, Okinawa 907-0002, Japan; Faculty of Life and Environmental Sciences, Shimane University, 1060 Nishikawatsu-cho, Matsue, Shimane 690-8504, Japan; Faculty of Applied Biological Sciences, Gifu University, 1-1 Yanagido, Gifu, Gifu 501-1193, Japan; National Institute for Agro-Environmental Sciences (NIAES), NARO, 3-1-3 Kannondai, Tsukuba, 305-8604, Japan

**Keywords:** early-morning flowering, flower opening time, heat stress, percentage of filled grains, rice, yield-related traits

## Abstract

Early-morning flowering (EMF) trait is supposed to be effective in retaining grain yield due to mitigation of heat-induced spikelet sterility at flowering in rice. This study evaluated (i) phenotypic differences between a near-isogenic line carrying a QTL for EMF trait, designated as IR64+*qEMF3*, and a recurrent parent, IR64, under wide variation in climates and (ii) whether an EMF trait can retain grain yield under heat stress at flowering. IR64+*qEMF3* had significant earlier flower opening time (FOT) in diverse environmental conditions including temperate, subtropical, and tropical regions. Under normal temperatures at flowering, IR64+*qEMF3* had similar grain yield to IR64 with some significant changes in agronomic traits and yield components. Field trials in heat-vulnerable regions of central Myanmar for seven crop seasons showed that higher percentage of filled grains contributed to the significantly higher grain yield in IR64+*qEMF3* among yield components when plants were exposed to daily maximum air temperatures around 36.5 °C or higher. Lower spikelet sterility in IR64+*qEMF3* was attributed to the earlier FOT during cooler early morning hours. This is the first field study that clearly demonstrates the advantage of the EMF trait for retaining grain yield by stabilizing percentage of fertile grains under heat stress at flowering.

## Introduction

Rice (*Oryza sativa* L.) is one of the most important staples feeding half the world’s population (Carriger and Vallee, 2007). While there is a need to produce more rice in order to feed the increasing population, challenges for increasing rice production have been rising due to progressive global warming (IPCC, 2013). Consequently, development of climate-smart rice cultivars is a pressing issue today. Genetic improvement of heat-resilience is one of the key requirements for enhancing global food security to tackle the increasing episodes of heat stress damage on crop production.

Major cereal crops show extreme sensitivity to heat during anthesis (Prasad et al., 2017). Chamber experiments conducted previously revealed that high proportion of sterility occurs in rice when spikelets are exposed to temperatures around 35 °C (Matsui et al. 1997; Satake and Yoshida, 1978) even for a short period of time (Jagadish et al., 2007). Actual increase in spikelet sterility by heat stress at flowering was reported in hot dry seasons of tropical and subtropical regions (Ishimaru et al., 2016; Matsushima et al. 1982; Osada et al., 1973). Rice crop is considered to be already at critical limits of heat stress vulnerability in central Thailand and Myanmar in dry season (Wassmann et al., 2009). Prediction models have identified high risk areas for reduction in rice grain yield by global warming in many rice-growing regions of Asia, mostly due to heat-induced spikelet sterility (See review, Horie, 2019). Central Myanmar has been projected as one among the heat-vulnerable regions for spikelet sterility and yield reduction in the future (Horie, 2019; Wassmann et al., 2009).

The EMF trait is proposed to be effective in escaping heat-induced spikelet sterility at flowering by shifting FOT to the cooler early morning times (Satake and Yoshida, 1978). Flower opening of modern cultivars predominantly occurs between 3.5–5.5 hours after dawn in the natural conditions of tropics (Bheemanahalli et al., 2017; Hirabayashi et al., 2015). Under elevated high temperature greenhouse conditions, spikelets that flower at later hours tend to be sterile with higher proportion; hence, shifting flower opening time to the cooler early morning, even a one-hour advancement, is effective in mitigating heat-induced spikelet sterility at flowering (Ishimaru et al., 2010). We previously developed a near-isogenic line (NIL) carrying a QTL for EMF (*qEMF3*) derived from a wild rice, *O. officinalis*, in the genetic backgrounds of IR64 (Hirabayashi et al., 2015). The NIL was designated as IR64+*qEMF3. qEMF3* advanced FOT by approximately 1.5–2.0 h in the tropical Philippines during both wet and dry seasons (Hirabayashi et al., 2015). Another field experiment also showed earlier FOT and lower sterility in IR64+*qEMF3* compared to a genetically diverse panel of germplasm in the dry rice growing season in Tamil Nadu, India (Bheemanahalli et al., 2017). However, whether the EMF trait has yield advantage under different heat stress levels on the field during flowering is yet to be ascertained. Whether *qEMF3* itself influences grain yield and other phenotypes of IR64 under normal temperatures during flowering stage is still not clear.

This study consists of two major objectives: the first is to evaluate advancement of FOT, agronomic traits, and yield performance of IR64+*qEMF3* in a wide range of field environments, including temperate, subtropical, and tropical regions, under normal rice-growing temperatures at flowering, and the second objective is to investigate if the EMF trait in the field helps in retaining rice grain yield in the heat-vulnerable region of central Myanmar.

## Materials and Methods

### Crop establishment

IR64, an Indica group cultivar, and a BC_3_-derived NIL with IR64 genetic background, IR64+*qEMF3* (Hirabayashi et al., 2015), were used for all six experimental sites, which were located in the temperate (Joetsu, Niigata, Japan), subtropical (Ishigaki, Okinawa, Japan), and tropical (Nay Pyi Taw, Myanmar; Los Baños, Laguna, Philippines; Coimbatore, Tamil Nadu, India; Subang, West Java, Indonesia) regions across five countries (Table 1), following the local rice cultivation practices (Supplementary Table S1). Plants were grown in fields during the summer/dry season (Table 1). Note that two staggered sowings were employed in Nay Pyi Taw for 2017DS, 2018DS, and 2019DS to expose rice plants to different temperatures at heading. 50% heading date was recorded for each genotype in each trial to determine the days-to-heading. Plant materials were kept healthy by spraying agrochemicals, when required.

**Table 1.** Geographical location of experimental sites and growth season(s).

| Experimental site | Geographical location | Above sea level (m) | Season(s) of experiment (summer/dry season) | Average daily maximum temperature during heading (°C)<sup>b</sup> |  |
| --- | --- | --- | --- | --- | --- |
|  |  |  |  | IR64 | IR64+*qEMF3* |
| CARC/NARO, Joetsu, Niigata, Japan | 37°1'N, 138°2'E | 10 | 2016 | 29.5 | 30.8 |
| JIRCAS, Ishigaki, Okinawa, Japan | 24°4'N, 124°2'E | 27 | 2015 | 32.8 | 33.1 |
| YAU, Nay Pyi Taw, Myanmar | 19°8'N, 96°3'E | 98 | 2015, 2017<sup>a</sup>, 2018<sup>a</sup>, 2019<sup>a</sup> | 35.0-37.9<sup>c</sup> | 35.0-38.3<sup>c</sup> |
| IRRI, Los Baños, Laguna, Philippines | 14°1'N, 121°2'E | 22 | 2014 | 29.6 | 27.7 |
| TNAU, Coimbatore, Tamil Nadu, India | 11°0'N, 76°5'E | 426 | 2020 | 34.4 | 35.5 |
| ICRR, Subang, West Java, Indonesia | 6°4'S, 107°6'E | 26 | 2015 | 32.2 | 32.2 |

<sup>a</sup>Two staggered sowings were employed in each dry season.
<sup>b</sup>Average daily maximum temperature during heading was calculated for six days around 50% heading date. 50% heading date was positioned as third day of six days.
<sup>c</sup>Average daily maximum temperature during heading in each season and genotype is indicated in Table 5.

### Flower opening time, spikelet sterility, and monitoring of local air temperatures

Flowering pattern observation and data analysis for the time to reach the peak and end of flower opening (FOT50 and FOT90, respectively) was conducted following the method of Hirabayashi et al., (2015). Using four heading panicles per plot per day, opened spikelets were marked every 30 min by fine-tipped pens, and this flowering observation was continued for more than three days in all sites. FOT50 and FOT90 were calculated using the Koyomi Station software (http://eco.mtk.nao.ac.jp/koyomi/index.html.en) and R program (ver. 3.6.2). Note that FOT50 and FOT 90 were presented as time after dawn. On 22 April of 2018, on Crop 1 in Nay Pyi Taw, opened spikelets were marked with different colored pens following the protocol of Ishimaru et al. (2010); spikelets opened before 0700H were marked with blue pens, those between 0700H-0830H with green pens, those between 0830H-1000H with purple pens, and those after 1000H with black pens. At maturity, marked spikelets marked with different colors were collected and sterility was physically checked. Air temperature during flowering was monitored by the meteorological station installed in each experimental site.

### Agronomic traits, grain yield, and yield components

At maturity, agronomic traits such as plant height (from ground level to the tip of panicles), panicle length (from panicle neck to the tip of panicle), flag leaf length, and flag leaf width were measured using the longest tiller in a hill. Grain yield and yield components per plant were determined in Joetsu, Nay Pyi Taw, Los Baños, Coimbatore, and Subang. Note that only one replicate was designed in Subang, but 3–5 replicates in other sites (Supplementary Table S1). Subsamples were prepared for selecting filled grains by an air-blower or water method and for subsequent determination of one thousand grain weight (1000GW). Selection of filled grains for seven yield trials in Nay Pyi Taw was consistently conducted by water method. Grain yield and 1000 GW were expressed based on 14% moisture content. Grain length, width, and thickness were measured with a digital caliper (SINWA Sokutei, Sanjo, Niigata, Japan) using the grains harvested in Joetsu, Ishigaki, and Nay Pyi Taw.

### Statistical analysis

Analysis of variance was tested by R program (ver. 3.6.2). Multiple regression analysis and *t-*test between genotypes were conducted by Microsoft Excel 2016.

## Results

### Flower opening time

Flowering pattern observation was conducted in Joetsu, Ishigaki, Nay Pyi Taw, Los Baños, and Subang under the condition that there was great variation in average daily maximum temperature during heading (Table 1; Supplementary Table 2). In Joetsu, Ishigaki, Los Baños, and Subang, FOT50 was significantly earlier in IR64+*qEMF3* than IR64 in each of the tested sites, ranging from 1.9 h to 3.5 h in IR64+*qEMF3* and 3.7 h to 4.6 h in IR64 on the basis of time after dawn (Table 2). FOT90 was also significantly earlier in IR64+*qEMF3* than in IR64, ranging from 2.5 h to 4.0 h in IR64+*qEMF3* and 4.2 h to 5.0 h in IR64 (Table 2). The genotype difference in the average of FOT50 and of FOT90 across locations was –1.5 h and –1.2 h, respectively (Table 2). In Nay Pyi Taw, flowering pattern observation conducted under a wide range of average daily maximum temperatures for seven crop seasons also showed significantly earlier FOT in IR64+*qEMF3* than IR64 (Supplementary Table 2).

**Table 2.**
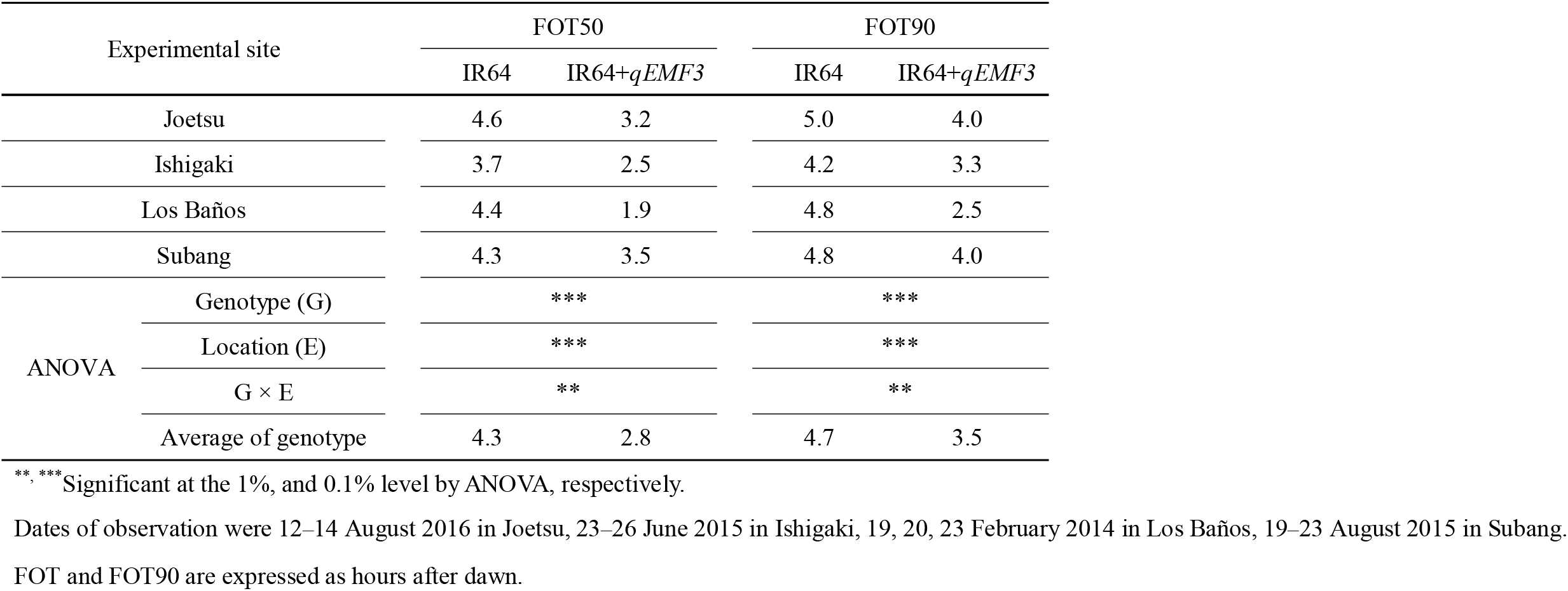
Time to reach the peak (FOT50) and end (FOT90) of flower opening in Joetsu, Ishigaki, Los Baños, and Subang.

### Agronomic traits, grain yield, and yield components under normal temperatures at flowering

Days-to-heading in IR64+*qEMF3* was the same as IR64 or up to five days longer than in IR64 (Tables 3, 4; Supplementary Table S3). Panicle length were not significantly different, while flag leaf length and flag leaf width were slightly larger in IR64+*qEMF3* (Table 3; Supplementary Table S3). Plant height was significantly higher in IR64+*qEMF3* than IR64 in Subang (Supplementary Table S3).

**Table 3.** Agronomic traits of IR64 and IR64+*qEMF3* in Joetsu, Nay Pyi Taw, Los Baños, and Coimbatore.

| Experimental site |  | Days to heading |  | Plant height (cm) |  | Panicle length (cm) |  | Flag leaf length (cm) |  | Flag leaf width (cm) |  |
| --- | --- | --- | --- | --- | --- | --- | --- | --- | --- | --- | --- |
|  |  | IR64 | IR64<br>*+qEMF3* | IR64 | IR64<br>*+qEMF3* | IR64 | IR64<br>*+qEMF3* | IR64 | IR64<br>*+qEMF3* | IR64 | IR64<br>*+qEMF3* |
| Joetsu |  | 109 | 111<sup>b</sup> | 99.3 | 99.9 | 23.9 | 23.2 | 21.7 | 21.0 | 1.25 | 1.31 |
| Nay Pyi Taw<sup>a</sup> |  | 101 | 101<sup>b</sup> | 84.8 | 83.8 | 23.5 | 22.8 | 20.5 | 21.1 | 1.25 | 1.28 |
| Los Baños |  | 82.6 | 86.4 | 83.3 | 82.4 | 22.8 | 22.1 | 29.4 | 30.8 | 1.24 | 1.28 |
| Coimbatore |  | 88.3 | 88.7 | 85.2 | 85.7 | 23.3 | 24.1 | 23.1 | 24.4 | 1.06 | 1.11 |
| ANOVA | Genotype (G) | - |  | ns |  | ns |  | † |  | ** |  |
|  | Location (E) | - |  | *** |  | *** |  | *** |  | *** |  |
|  | G × E | - |  | ns |  | † |  | ns |  | ns |  |
|  | Average of genotype | 95.2 | 96.8 | 88.2 | 88.0 | 23.4 | 23.1 | 23.7 | 24.3 | 1.20 | 1.25 |

†, \*\*, \*\*\* Significant at the 10%, 1%, and 0.1% level, respectively, by ANOVA; ns: not significant.
<sup>a</sup>Measurement in Nay Pyi Taw was conducted in 2015DS.
<sup>b</sup>Days-to-heading data was taken only from one plot with intermediate days-to-heading for each genotype.

**Table 4.**
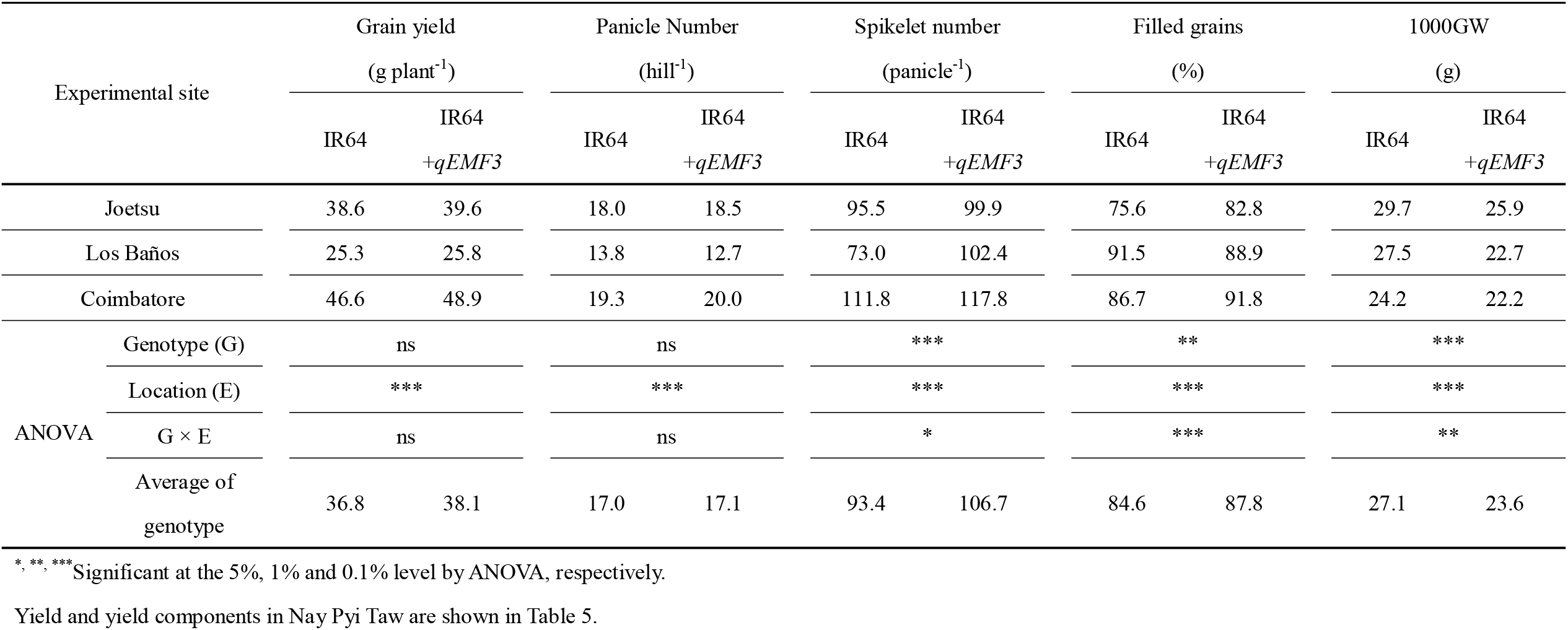
Grain yield and yield components in IR64 and IR64+*qEMF3* at normal temperatures during heading.

### Grain yield and yield components under normal temperatures at flowering

Grain yield and yield components were further examined in Joetsu, Los Baños, Coimbatore, and Subang (Table 4; Supplementary Table S3), where average daily maximum temperatures at flowering was in the normal ranges below 35.5 ^°^C (Table 1). Grain yield and panicle number per plant (PN) was not significantly different between IR64 and IR64+*qEMF3*, but spikelet number per panicle (SN) was significantly larger and 1000GW was significantly smaller in IR64+*qEMF3* (Table 4; Supplementary Table S3). Smaller 1000GW in IR64+*qEMF3* was mainly due to the smaller length and width of grains compared to IR64 grains (Supplementary Table S4). Percentages of filled grains (PFG) was similar or slightly higher in IR64+*qEMF3* on an average, but the trend was not the same on all locations (Table 4; Supplementary Table S3).

### Grain yield and yield components in heat-vulnerable regions of central Myanmar

Grain yield and yield components were evaluated in heat-vulnerable regions of central Myanmar for seven crop seasons across four years (Table 5). Average daily maximum temperatures during heading ranged from 35.0 ^°^C to 38.3 ^°^C (Table 5). Grain yield per plant was significantly higher in IR64+*qEMF3* than IR64, resulting in yield advantage of IR64+*qEMF3* by 22.4% over IR64 on an average. PN was similar between genotypes, while SN and 1000GW was significantly lower in IR64+*qEMF3* than in IR64. Such a trend in PN, SN, and 1000GW characters was similar in other locations as well, as shown in Table 4. Significant interaction between the genotype and environment was observed in PFG; difference in PFG between genotypes in each season ranged from 0.8% to 17.4%, and PFG was 9.3% higher in IR64+*qEMF3* than in IR64 on an average (Table 5). Multiple regression analysis revealed a significant positive effect of PFG on yield advantage of IR64+*qEMF3*, while other parameters such as PN, SN and 1000GW did not have a significant effect (Figure 1). The relationship between PFG and average daily maximum temperatures during heading is plotted in Figure 2. Difference in PFG between genotypes was small when average daily maximum temperatures during heading was below 36.5 ^°^C. PFG steadily decreased in IR64 when average daily maximum temperatures during heading was greater than 36.5 ^°^C, and it sharply dropped to 65.6% at 37.9 ^°^C (Figure 2). On the contrary, PFG was stable at the high level of around 90% until 37 ^°^C and gradually decreased to 83.0% at 38.3 ^°^C in IR64+*qEMF3* (Figure 2).

**Table 5.** Heading date, average of daily maximum temperature during heading, grain yield and yield components in IR64 and IR64+*qEMF3* in Nay Pyi Taw.

| Year | Crop | 50% heading date |  | Average daily maximum temperature during heading (°C) |  | Grain yield (g plant<sup>-1</sup>) |  | Panicle number (plant<sup>-1</sup>) |  | Spikelet number (panicle<sup>-1</sup>) |  | Filled grains (%) |  | 1000GW (g) |  |
| --- | --- | --- | --- | --- | --- | --- | --- | --- | --- | --- | --- | --- | --- | --- | --- |
|  |  | IR64 | IR64<br>+*qEMF3* | IR64 | IR64<br>+*qEMF3* | IR64 | IR64<br>+*qEMF3* | IR64 | IR64<br>+*qEMF3* | IR64 | IR64<br>+*qEMF3* | IR64 | IR64<br>+*qEMF3* | IR64 | IR64<br>+*qEMF3* |
| 2015 | Crop 1 | 3 April | 3 April | 35.5 | 35.5 | 24.9 | 28.5 | 16.6 | 16.9 | 67.3 | 84.5 | 90.8 | 91.6 | 24.7 | 21.8 |
| 2017 | Crop 1 | 1 May | 2 May | 37.9 | 38.3 | 17.6 | 25.0 | 14.0 | 14.3 | 79.1 | 96.8 | 65.6 | 83.0 | 24.7 | 21.1 |
| 2017 | Crop 2 | 18 May | 19 May | 37.1 | 37.6 | 25.0 | 29.7 | 16.4 | 14.5 | 104.2 | 116.0 | 75.0 | 85.8 | 23.6 | 20.8 |
| 2018 | Crop 1 | 21 April | 23 April | 36.6 | 36.5 | 21.8 | 26.5 | 13.8 | 14.0 | 78.3 | 89.0 | 83.4 | 94.1 | 24.2 | 22.6 |
| 2018 | Crop 2 | 18 May | 18 May | 35.0 | 35.0 | 22.8 | 24.7 | 13.1 | 12.5 | 81.3 | 85.7 | 86.1 | 91.3 | 25.1 | 25.3 |
| 2019 | Crop 1 | 5 April | 9 April | 36.4 | 36.9 | 25.4 | 31.5 | 18.0 | 19.6 | 62.9 | 77.9 | 85.0 | 92.9 | 27.2 | 22.9 |
| 2019 | Crop 2 | 4 May | 9 May | 36.9 | 37.9 | 37.6 | 48.2 | 28.2 | 30.2 | 76.7 | 83.0 | 76.2 | 88.2 | 23.5 | 22.2 |
| ANOVA |  | Genotype (G) |  | - | - | *** |  | ns |  | *** |  | *** |  | *** |  |
|  |  | Season (E) |  | - | - | *** |  | *** |  | *** |  | *** |  | *** |  |
|  |  | G × E |  | - | - | ns |  | ns |  | ns |  | ** |  | ns |  |
|  |  | Average of genotype |  | 36.5 | 36.8 | 25.0 | 30.6 | 17.2 | 17.4 | 78.5 | 90.4 | 80.3 | 89.6 | 24.7 | 22.4 |

\*\*, \*\*\* Significant at 1% and 0.1% level by ANOVA, respectively.

**Figure 1.**
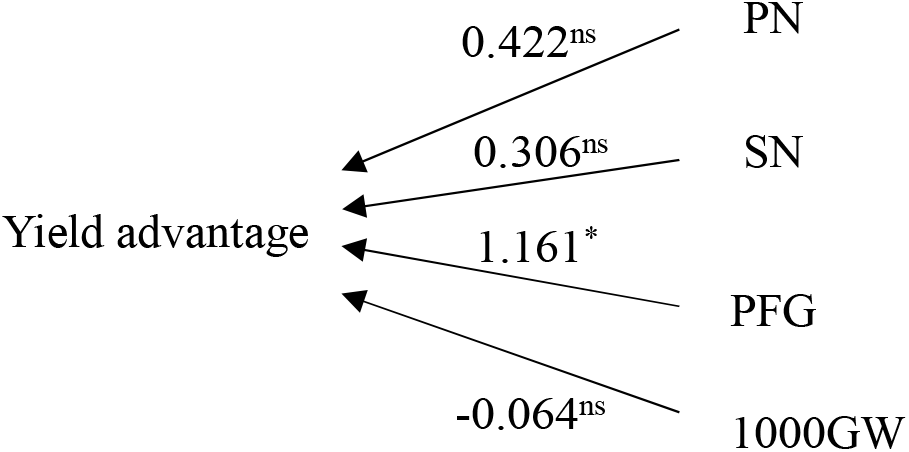
A diagram showing multiple regression of yield advantage in IR64+*qEMF3* on yield components, along with the correlation between explanatory variables for the data obtained in seven seasons of Nay Pyi Taw. Values with the single head arrows are standardized multiple regression coefficients. ^*^, P<0.05; ns, not significant.

**Figure 2.**
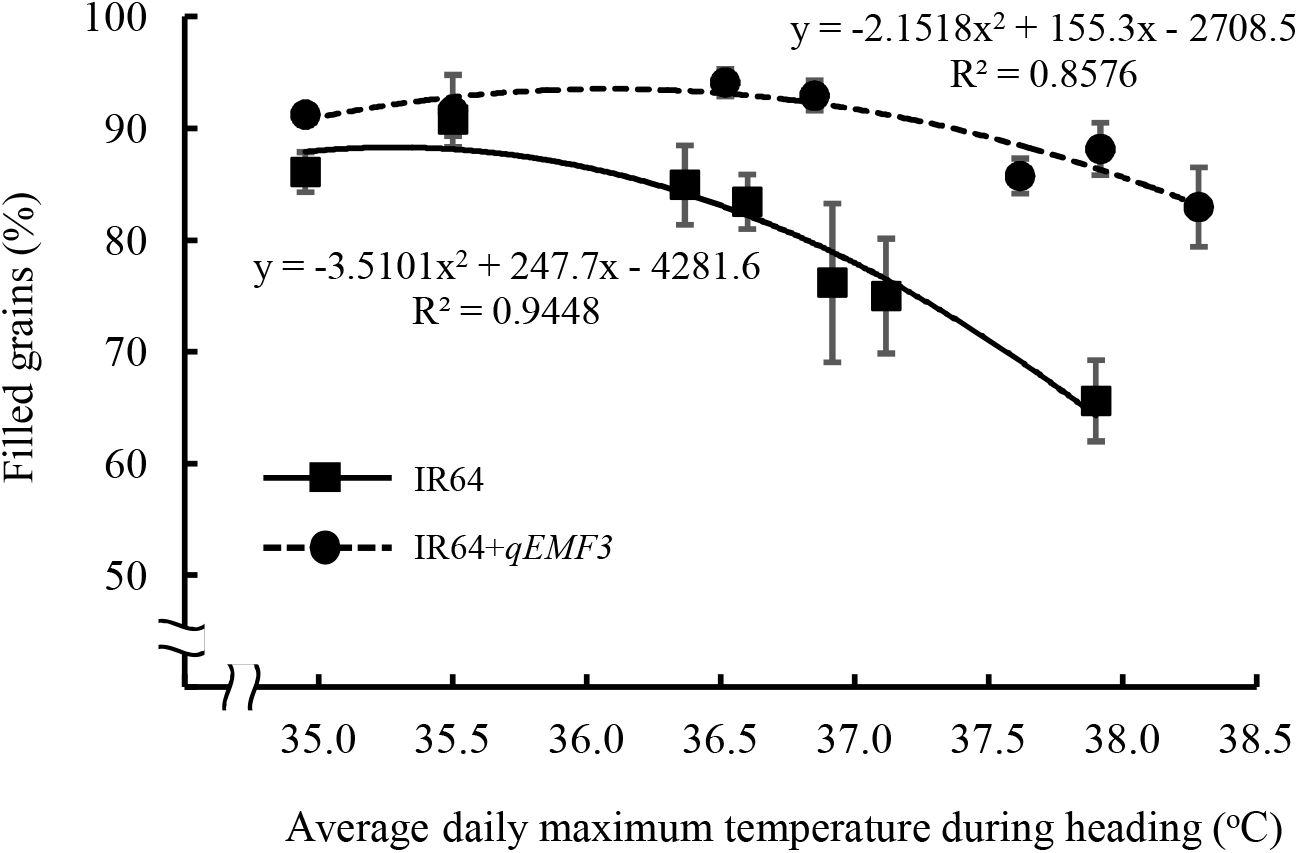
Relationship between average daily maximum temperatures during heading and percentages of filled grains. Bars indicate means of percentages of filled grains ± standard deviation.

Spikelet sterility was examined in different time periods of flower opening (Figure 3). During flowering pattern observation (recorded with different colored pens), air temperature steadily increased from 25.8 ^°^C at 0700H to 30.0 ^°^C at 0930H, and up to 34.0 °C at noon (Figure 3A). Under such air temperature condition, flower opening in IR64+*qEMF3* started after 0730H, peaked at 0900H, and almost ended by 0930H (Figure 3A). On the other hand, flower opening in IR64 started after 0830H, peaked at 0930H, and ended by 1030H (Figure 3A). Spikelet sterility in each time period was 3.2% and 8.6% for spikelets that flowered during 0700-0830H (green-lined bar) and during 0830-1000H (purple-lined bar), respectively, in IR64+*qEMF3* (Figure 3B). On the other hand, spikelet sterility was 10.2% and 47.2% for spikelets flowered during 0830-1000H (purple bar) and after 1000H (black bar), respectively, in IR64 (Figure 3B).

**Figure 3.**
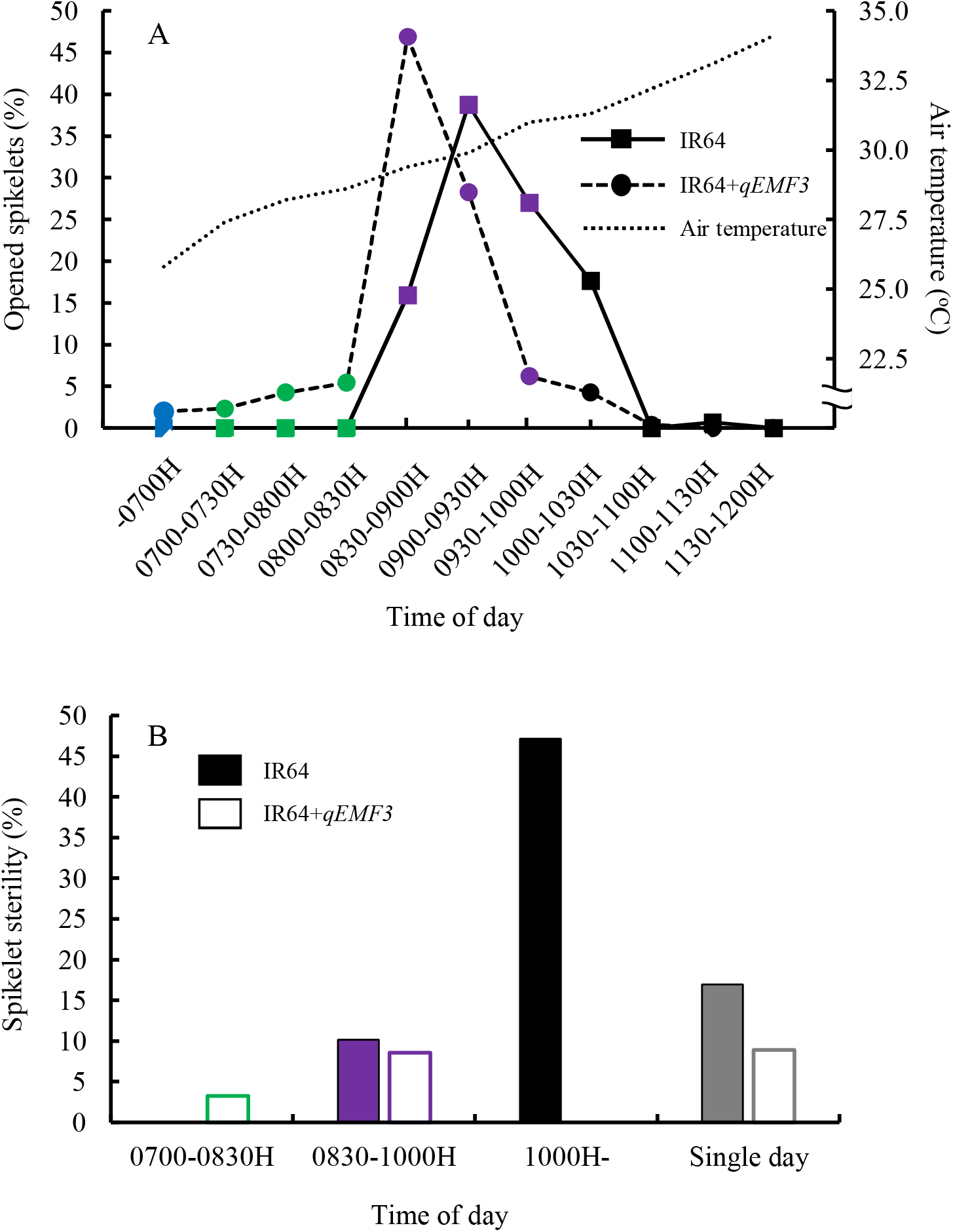
Daily flowering pattern and change in air temperature (A), and changes in percentages of spikelet sterility in each time period and on a single day (B) on 22 April 2018. Bars shown in blue, green, purple, and black colors indicate that the spikelets opened before 0700H, between 0700H-0830H, 0830H-1000H, and after 1000H, respectively. In B, color bars for IR64 and white bars with color outline for IR64+*qEMF3* are shown.

As a consequence, spikelet sterility was 17.0% and 8.9% in IR64 and IR64+*qEMF3*, respectively, on that single day (Figure 3B).

## Discussion

### Significant advancement of FOT by qEMF3 in a wide range of environmental conditions

Hirabayashi et al., (2015) and Bheemanahalli et al., (2017) previously reported an earlier FOT in IR64+*qEMF3* than IR64 in Philippines (dry and wet seasons) and southern India (dry season), under field conditions. This study validated the genetic gain in advancing FOT by *qEMF3* under diverse environments including temperate to tropical climates in the genetic background of IR64 (Table 2; Supplementary Table S2). It should be noted, however, that the difference in FOT between IR64 and IR64+*qEMF3* varies depending on the experimental site or seasons tested (Table 2; Supplementary Table S2). It is possible that this difference is because FOT is determined in the fields by microclimates such as solar radiation and vapor pressure deficit in addition to air temperature (Julia and Dingkuhn, 2012; Kobayasi et al., 2010). Development of a prediction model for FOT that takes into consideration various such meteorological parameters will help us in identifying the locations where *qEMF3* can make a significant advancement in FOT to escape heat stress at flowering. The nature of interactions between genetic and environmental factors on FOT in IR64+*qEMF3* is still not very clear.

### qEMF3 helps retain grain yield by stabilizing percentage of filled grains under heat stress at flowering

Central Myanmar is projected as one of the heat-vulnerable regions for spikelet sterility and yield reduction in the future (Horie, 2019; Wassmann et al., 2009). By employing staggered sowings in different dry seasons, wide range of average daily maximum temperatures during heading was obtained (Table 5). IR64+*qEMF3* was tested under such field conditions to examine if an EMF trait could mitigate the yield loss under heat stress at flowering.

PFG was similar between genotypes or slightly higher in IR64+*qEMF3* under normal temperature at flowering (Table 4; Supplementary Table S3), whereas a total of seven trials in the central Myanmar revealed that PFG greatly varied depending on the seasons (Table 5). Genetic and environmental interaction were not observed in other yield components such as PN, SN, and 1000GW (Table 5). A clear difference in response of PFG to average daily maximum temperature at heading between genotypes was observed (Figure 2). PFG steadily dropped in IR64 as average daily maximum temperature at heading increased above 36.5 ^°^C (Figure 2). On the other hand, high PFG of greater than 80% was maintained in IR64+*qEMF3* even when average daily maximum temperature at heading was around 38 ^°^C (Figure 2). Among yield components, PFG was the only parameter that significantly contributed to the yield advantage of IR64+*qEMF3* (Figure 1). Flowering pattern observation revealed less frequency of spikelet sterility in IR64+*qEMF3* than in IR64 on a single day due to the earlier FOT in IR64+*qEMF3* (Figure 3A, 3B). These results indicate the yield advantage that *qEMF3* conferred under high temperature conditions at heading by significant advancement of FOT in IR64 genetic background. Notably, an approximately one-hour difference in FOT50 and FOT 90 in Crop 1 of 2018 (Supplementary Table S2) made a difference in spikelet sterility between IR64 and IR64+*qEMF3* (Figure 3B), supporting that one-hour advancement of FOT is sufficient to avoid heat-induced spikelet sterility as shown in previous chamber experiments (Ishimaru et al. 2010; Satake and Yoshida 1978).

Horie (2019) documented that EMF is one of the key traits that could retain rice grain yield by mitigating heat-induced spikelet sterility during flowering under future hotter temperatures in inland of continental South East Asia. Our study clearly demonstrated that EMF trait could diminish heat stress damage at flowering on grain yield through stabilization of PFG in the hot dry season of central Myanmar. IR64 is a moderately heat-tolerant cultivar (Shi et al., 2015). Some of Myanmar cultivars could be reaching critical limits of heat tolerance as described by Wassmann et al., (2009). Assessment of heat tolerance among Myanmar local cultivars helps to identify the target cultivars to be conferred with EMF trait. Heat-vulnerable regions, in terms of heat-induced spikelet sterility, are estimated to be located in different geographic locations including China, South-East Asia, South Asia, and West Africa (Laborte et al., 2012). Since IR64+*qEMF3* can significantly advance FOT in diverse environmental conditions (Table 2; Supplementary Table S2), *qEMF3* may potentially minimize reduction in heat-induced spikelet sterility across regions. It should be noted, however, microclimate including humidity (Matsui et al., 2014; Tian et al., 2011), wind velocity (Ishimaru et al., 2012; Matsui et al., 1997), and solar radiation (Ishimaru et al., 2016), which are not inclusive in analysis in the present study, have a complex interaction effect on spikelet sterility under field heat stress through changes in panicle temperature (Yoshimoto et al., 2011). Whether IR64+*qEMF3* can mitigate yield loss in other heat-vulnerable regions with different microclimate needs to be further tested.

### Further breeding efforts for the genetic improvement of IR64+qEMF3 and other backgrounds

Some yield components were significantly different between genotypes. SN tended to be higher while 1000GW was lower in IR64+*qEMF3* than in IR64 (Table 4, 5; Supplementary Table S3). However, PN was not different between genotypes (Table 4; Supplementary Table S3). IR64+*qEMF3* had larger flag leaf size compared to IR64 (Table 3; Supplementary Table S3). Smaller 1000GW in IR64+*qEMF3* was mainly due to the smaller length of the grains (Supplementary Table S4). These results suggest that physical constraints due to small husk size, and not insufficient source supply, would be responsible for the smaller 1000GW observed. Ohsumi et al., (2011) demonstrated that there is a negative correlation between spikelet number per panicles and 1000GW using NILs for increased spikelet number. Whether *qEMF3* has pleiotropic effects on SN and 1000GW as well as the EMF trait must be further investigated. Introgression of QTL for grain size detected in genetic background of Indica group cultivars (Kato et al., 2011; Qing et al., 2018) may retrieve 1000 GW of IR64+*qEMF3* to similar levels as those of IR64. Regarding other agronomic traits, minor change in days-to-heading were observed (Table 3, 5; Supplementary Table S3). IR64 is a popular Indica group cultivar that is most widely grown in Asia (Brennan and Malabayabas, 2011) and its progeny continues to be grown quite widely. Our study provides breeders some fundamental information on the changes in plant phenotypes that can occur due to the introgression of *qEMF3* to IR64 genetic background. Transfer of *qEMF3* to popular cultivars in the temperate, subtropical, and tropical regions by marker-assisted breeding is an ongoing project. The effects of *qEMF3* on agronomic traits and grain yield on other genetic backgrounds need to be further investigated.

## Conclusion

This is the first field study to clearly demonstrate that the EMF trait can retain rice grain yield through stabilization of PFG under stressful conditions in heat-vulnerable regions. Significant advancement of FOT due to *qEMF3* was observed in wide range of environments including temperate, subtropical, and tropical regions, while the changes in agronomic traits due to the introgression of *qEMF3* into genetic background of IR64 were minimal. This result indicates the possibility that *qEMF3* can also be effective in retaining rice grain yield in other heat-vulnerable regions and genetic backgrounds. A breeding program to transfer *qEMF3* to local cultivars grown in hot dry seasons through marker-assisted selection is an appropriate strategy that is recommended.

## Supplementary data

Supplementary Table S1 Experimental design and fertilizer conditions in each experimental site.

^a^ The experimental plots were arranged in a randomized complete block design in Joetsu, Los Baños, Nay Pyi Taw, and Coimbatore.

^b^ 60 panicles that were randomly chosen from the bulk of panicles were used for the determination of yield and yield components.

Supplementary Table S2 Time to reach the peak (FOT50) and end (FOT90) of flower opening in Nay Pyi Taw.

^***^Significant at 0.1% level by ANOVA.

Dates of observation were 3–6 April in Crop 1 of 2015, 28–30 April and 1 May in Crop 1 of 2017, 16–19 May in Crop 2 of 2017, 21–24 April in Crop 1 of 2018, 16–19 May 12–14 in Crop 2 of 2018, 4–7 April for IR64 and 8–11 April for IR64+*qEMF3* in Crop 1 of 2019, and 3–6 May for IR64 and 8–11 May for IR64+*qEMF3* in Crop 2 of 2019.

FOT50 and FOT90 are expressed as hours after dawn. Average of maximum temperature during heading is shown in Table 5.

Supplementary Table S3 Agronomic traits (upper), yield, and yield components (lower) of IR64 and IR64+*qEMF3* in Subang.

Measurement was conducted by individual 14 plants except for Days to heading.

^†, **, ***^Significant at the 10%, 1%, and 0.1% level, respectively, by *t*-test.

Supplementary Table S4 Physical dimensions of the grain in IR64 and IR64+*qEMF3*.

^a^ Measurement in Nay Pyi Taw was done using grains harvested in Crop 1 of 2019.

## Acknowledgements

We thank research technicians who were in charge of field management in each experimental site. Meteorological data in Ishigaki and Los Baños was provided by Dr. S. Goto and Dr. K. Okamoto (TARF/JIRCAS), and IRRI Climate Unit, respectively. T.I. thanks Dr. K.S.V. Jagadish (Kansas State University) and Dr. M. Okamura for critical reading of the manuscript and advice in statistical analysis, respectively. This work was supported by the Japanese government under the IRRI-Japan Collaborative Research Project (to T. I.), by grant-in-aid from the Ministry of Education, Culture, Sports, Science and Technology of Japan (No.26257410 to T. M., K. K., and T. I.), and Global Environment Research Coordination System organized by Ministry of Environment (To M. Y. and T. I.).

## Abbreviations

CARC/NARO: Central Region Agricultural Research Center, National Agriculture and Food Research Organization
DAR: Department of Agricultural Research
EMF: early-morning flowering
FOT: flower opening time
ICRR: Indonesian Center for Rice Research
JIRCAS: Japan International Research Center for Agricultural Sciences
NIL: near-isogenic line
PFG: percentage of filled grains
PN: panicle number per hill
SN: spikelet number per panicle
TNAU: Tamil Nadu Agricultural University
YAU: Yezin Agricultural University
1000GW: one thousand grain weight

